# Data rescue in high-motion youth cohorts for robust and reproducible brain-behavior relationships

**DOI:** 10.1101/2024.06.04.597447

**Authors:** Jivesh Ramduny, Tamara Vanderwal, Clare Kelly

**Author notes:** **CORRESPONDING AUTHOR:** Clare Kelly, Trinity College Institute of Neuroscience, Trinity College, Dublin 2, Ireland.

## Abstract

Recognition of the detrimental impact of participant motion on functional connectivity measures has led to the adoption of increasingly stringent quality control standards to minimize potential motion artifacts. These stringent standards can lead to the exclusion of many participants, creating a tension with a countervailing requirement for large sample sizes that can provide adequate statistical power, particularly for brain-behavior association studies. Here, we test and validate two techniques aimed at mitigating the impact of head motion on functional connectivity estimates, and show that these techniques enable the retention of a substantial proportion of participants who would otherwise be excluded based on motion criteria, such as a minimum mean framewise displacement (FD) threshold. Specifically, we first show that functional connectomes computed using time series data that have been ordered according to motion (i.e., framewise displacement — FD) and (1) subsetted to include the lowest-motion time points (“*motion ordered*”) or (2) subsetted and resampled (“*bagged*”) are reproducible, in that they enable the successful identification of an individual from a group using functional connectome fingerprinting. Second, we demonstrate that motion-ordered and bagged functional connectomes yield robust brain-behavior associations, which, when examined as a function of sample size, are comparable to those obtained using the standard full time series. Finally, we show that the utility of both approaches lies in maximizing participant inclusivity by allowing for the retention of high-motion participants that would otherwise be discarded. Given equivalent performance of the two approaches across these tests of reproducibility, validity, and utility, we conclude by recommending motion-ordering to enable data rescue, maximize inclusivity, and address the need for adequately powered samples in functional connectivity research, while maintaining stringent data quality standards. While our findings were reproducible across different head motion thresholds and edges in the functional connectome, we outline possibilities for further validation and assessment of generalization using other behavioral phenotypes and consortia datasets.

## 1 Introduction

Careful analyses of the effects of participant head motion during scanning on fMRI-based measures of functional connectivity and of the impact of inadequate sample size and low statistical power on the robustness and reproducibility of findings obtained using such measures (Szucs and Ioannidis, 2020; Cremers et al., 2017; Button et al., 2013; Thirion et al., 2007) have prompted a beneficial shift by the neuroimaging community towards both retro- and prospective data sharing and pooling of fMRI data from multiple sites (Casey et al., 2018; Alexander et al., 2017; Miller et al., 2016; Zuo et al., 2014; Van Essen et al., 2013; Di Martino et al., 2014b; Biswal et al., 2010). These efforts better enable the application of strict data quality standards, to mitigate the confounding influence of motion-related and other artifacts, while meeting sample size demands, to provide adequate statistical power for reproducible brain-behavior associations.

Nevertheless, these two countervailing imperatives can create a tension between sample size and data quality by discarding participants whose data are perceived to be “unusable” for example, due to excess in-scanner motion, we sacrifice sample size in favor of quality. In the context of very large consortia datasets spanning to thousands of participants, this trade-off may appear manageable. However, it limits the diversity of the samples included in our brain-behavior analyses. Participants who are excluded by the application of strict motion thresholds are those who are the youngest and/or who experience more severe clinical conditions (Nebel et al., 2022; Frew et al., 2022; Vanderwal et al., 2019; Engelhardt et al., 2017; Vanderwal et al., 2015; Wylie et al., 2014; Durston et al., 2003). Excluding those participants truncates the range of the symptom or behavioral dimension under examination. In the context of brain-behavior analyses, this directly impedes reproducibility — range restriction violates the assumptions of statistical analyses such as correlation by reducing the representativeness of the sample, relative to the population (Goodwin and Leech, 2006). Restricting the range of symptoms and/or behaviors also reduces the effect sizes of brain-behavior relationships by decreasing the variability of the data (Gratton et al., 2022).

Instead of discarding participants whose in-scanner motion exceeds a given threshold, here, we propose two methods to “rescue” these individuals’ data so that they can be retained in analyses, thus boosting sample size and diversity for the examination of brain-behavior associations. The first is a variant of the motion-ordered approach proposed by Fair et al. (2020). In their paper, Fair et al. (2020) ordered participant time series according to head motion (measured by framewise displacement; FD). Functional connectivity matrices were then computed using a sliding window with a length of 150 frames, and a mean FD-filtered functional connectivity matrix was created by averaging the functional connectivity matrices from the first 30 windows (i.e., those that included the time points with the lowest FD values). The authors showed that the fMRI data quality and reliability of functional connectivity measurements were improved using this approach. Here, we first “scrubbed” each participants’ time series by removing time points with root mean square FD (rmsFD) > 0.20 mm, the preceding time point (*T* – 1) and two succeeding time points (*T* + 1, *T* + 2) (Power et al., 2015; Power et al., 2014; Power et al., 2013). Next, following Fair et al. (2020), we ordered each individual’s time series according to head motion (indexed by rmsFD). We then retained a fixed number of time points with the least motion (e.g., the 100 least motion-contaminated time points). We then computed functional connectivity as normal; we refer to this as the motion-ordered approach.

Next, instead of applying the sliding window approach described by Fair et al. (2020) (since this requires at least 180 time points), we implemented an approach known as “bagging” or bootstrap aggregation, at the time series level (Opitz and Maclin, 1999; Breiman, 1996). In its typical application in machine learning contexts, bagging means resampling *with* replacement to create multiple random subsets of training data. These subsamples are then used to generate several instances of a machine learning model and their predictions are aggregated to yield more robust performance estimates. A main advantage of this bootstrap sampling technique is that it can minimize variability and bias in a noisy high-dimensional dataset (Varoquaux et al., 2017; Dudoit and Fridlyand, 2003) by reducing model underfitting (i.e., low variance and high bias) and/or overfitting (i.e., high variance and low bias), in addition to improving model generalizability in unseen datasets (Varoquaux et al., 2017; Dudoit and Fridlyand, 2003). Bagging can be applied at the level of samples by computing bootstrap subsets of the training data or at the level of time series themselves by sampling random time points — both across a number of iterations (e.g., ≥ 100).

An approach similar to bagging, jack-knife resampling, was first applied to fMRI data in a proof-of-concept study to estimate confidence intervals (CIs) for correlation coefficients indexing task-evoked activation during a finger-tapping paradigm (Biswal et al., 2001). Specifically, 85 time points were repeatedly resampled (*without* replacement) from a time series of 90 time points acquired during 4.5 cycles of 20 seconds alternating finger-tapping/rest. For each of the 1,000 jack-knife samples, the box-car reference waveform was also resampled in a matched fashion and the correlation between the reordered fMRI time series and waveform was computed. This process produced an approximation of the theoretical population from the observed data (sample), enabling the estimation of reliability and CIs for the parameter of interest (here, correlation with the reference waveform). Biswal et al. (2001) then generated maps of task-evoked activation thresholded on the basis of the estimated distributions, rather than a simple fixed correlation coefficient threshold. Biswal et al. (2001) argued that this approach produced more robust, sensitive activation maps because it better accounts for variation in both signal (response) and noise across voxels. They also suggested that their approach reduced the chance of false positives, thus increasing reproducibility.

Combining both approaches, Bellec et al. (2010) and Nikolaidis et al. (2020) used bagging *within* individual participants’ time series and *across* participants to improve the reproducibility of functional brain parcellations derived from relatively short acquisition scans (e.g., 6 min). Specifically, Nikolaidis et al. (2020) applied bagging to create multiple bootstrap resampled time series (using circular block resampling) for each participant. Clustering was applied to these resampled time series and cluster solutions were transformed to adjacency matrices, which were then averaged for each participant. Bagging was then applied again at the participant level — individual adjacency matrices were averaged within many resampled groups, and clustering was applied to these group-mean adjacency matrices to obtain a final cluster solution. This approach increased the test-retest reliability and reproducibility of functional connectivity-based parcellations across a number of acquisition (e.g., scan lengths, scanner, imaging sites) and processing (e.g., clustering parameters and algorithms) parameters. Moreover, reproducibility for the bagging approach applied to 6-min scans was greater than that for the standard parcellation approach applied to scans that were double the duration. This suggests that bagging may compensate for the effects of short scan durations — an important finding when considering the potential application of this method to time series that have been shortened due to the exclusion of time points that exceed head motion thresholds. In the present study, our focus is on bagging at the level of individual time series. Our bagging approach is conceptually similar to that used by Biswal et al. (2001), Bellec et al. (2010), and Nikolaidis et al. (2020), with a key difference being that, rather than randomly resampling time points or blocks of time points from an individual time series, we first sorted each individual’s scrubbed time series according to head motion and retained a fixed number of time points with the least motion (i.e., as in the motion-ordered approach described above). Next, we performed bootstrap resampling (*with* replacement) within this subset to create multiple bootstrap resampled time series for each participant. We computed functional connectivity for each bootstrapped sample and averaged across samples to derive an average bagged functional connectivity matrix for each participant.

In the present study, we assessed the reproducibility of the motion-ordering and bagging approaches by examining their effects on functional connectome fingerprinting (Finn et al., 2015). We complemented this test of reproducibility with an examination of validity — following Marek et al. (2022), we examined the impact of the motion-ordering and bagging approaches on estimates and CIs for brain-behavior relationships as a function of sample size. For both analyses, the results obtained using motion-ordered and bagged functional connectivity matrices were compared to those obtained using standard, scrubbed series. We hypothesized that the motion-ordering and bagging approaches would yield reproducible functional connectivity matrices and robust brain-behavior relationships, but expected bagging to exceed motion-ordering alone, in terms of performance. We show that both approaches equivalently enable the “rescue” of high-motion participants who would otherwise be excluded, and thus recommend the simpler motion-ordering approach as a strategy to enable researchers to maximize sample size and compute more inclusive and reproducible brain-behavior relationships.

## 2 Methods

### 2.1 Participants

Data were obtained from the Healthy Brain Network (HBN; (http://fcon_1000.projects.nitrc.org/indi/cmi_healthy_brain_network/), which is a community-based help-seeking sample of children and adolescents from the Greater New York area (Alexander et al., 2017). Neuroimaging and phenotypic data were obtained from the Cornell Brain Imaging Center (CBIC) site, Releases 4-9, comprising 682 participants. This dataset was selected as representative of a challenging dataset — including participants as young as 6 and enriched for clinical symptoms, such as internalizing and externalizing symptoms. Ethical approval for data collection and sharing was obtained from the Chesapeake Institutional Review Board; written consent was provided by all the participants who were at least 18 years old, while written consent was obtained from legal guardians and from the participants who were under 18 years old. The current study procedures were approved by the School of Psychology Research Ethics Committee, Trinity College Dublin. For each participant, at least one anatomical and two resting-state fMRI scans were obtained from the same imaging session (i.e., within-session scans). All participants were required to have two resting-state fMRI scans to enable functional connectome fingerprinting. Quality control was performed on all the scans to check for potential scan artifacts in addition to ensuring that each participant had two runs of resting-state fMRI data as well as their phenotypic data to reflect their age and sex. Note that motion was addressed as part of the analyses; no initial head motion threshold was applied. A total of 423 participants (153 females; age range = 6-20 years; mean age±SD = 10.7±3.0 years) remained in the cohort for subsequent analyses.

### 2.2 MRI Data Acquisition

Structural T1-weighted and functional scans were acquired at the CBIC using a Siemens 3T Prisma scanner with a 32-channel head coil. A 3D T1-weighted MPRAGE volume with spatial resolution 0.8 x 0.8 x 0.8 mm^3^ was obtained for each participant (224 sagittal slices, TR = 2500 ms, TE = 3.15 ms, flip angle = 8°). The two runs of resting-state fMRI data were acquired with echo-planar imaging sequences (60 slices, voxel size = 2.4 x 2.4 x 2.4 mm^3^, multiband factor = 6, TR = 800 ms, TE = 30 ms, flip angle = 31°). The participants underwent two 5 min functional runs while they fixated a cross on the screen and both runs consisted of 375 volumes.

### 2.3 MRI Preprocessing Pipeline

All data were preprocessed using a standard preprocessing pipeline (Yan et al., 2013a; Yan et al., 2013b; Craddock et al., 2013). Structural and functional data were preprocessed using AFNI (http://afni.nimh.nih.gov/afni/), FSL (http://fsl.fmrib.ox.ac.uk/), and ANTs (http://stnava.github.io/ANTs). The preprocessing pipeline included: (1) volume-based motion correction; (2) grand mean scaling; (3) linear and quadratic detrending; (4) nuisance signal regression on 24 motion parameters (i.e., 3 translational, 3 rotational, their first derivatives, and the squares of each of these terms) and 12 nuisance regressors (i.e., white matter, cerebrospinal fluid, global signal, their squares, and the squares of their first derivatives); (5) spatial smoothing using a Gaussian kernel at 6mm FWHM; and (6) temporal filtering (0.009 ≤ ƒ ≤ 0.1 Hz). Functional-to-anatomical co-registration was performed using boundary-based registration in FSL (Greve and Fischl, 2009). Diffeometric normalization of individual anatomical to the MNI152 standard space was carried out using ANTs; the same non-linear transformation was applied to the fMRI data.

### 2.4 Head Motion

To identify high-motion participants in the HBN dataset, a head motion threshold of root-mean-square framewise displacement [msFD] > 0.20 mm was applied (Satterthwaite et al., 2012; Jenkinson et al., 2002). After applying a rmsFD > 0.20 mm, a total of 255 participants remained in the sample which corresponds to ∼60% of the original sample size.

### 2.5 Parcellation Scheme

We used a whole-brain parcellation scheme to compute the functional connectivity matrices of each participant (https://neurovault.org/images/395091/; Shen et al., 2013). The Shen 268 parcellation scheme describes the cortical, subcortical, and cerebellar regions with a reasonable number of parcels. When the Shen 268 parcellation scheme was applied, a total of [(268 x 268) – 268] / 2 = 35,778 unique edges were obtained in the whole-brain functional connectome.

### 2.6 Functional Connectivity: Standard Approach

Using the Shen 268 parcellation scheme, the mean time series of each ROI was extracted prior to computing the Pearson correlation coefficient (*r*) between all possible ROI pairs to construct a 268 x 268 functional connectivity matrix for each participant. Hence, the correlation values represent the connectivity strengths (edges) between two ROIs (nodes). This procedure was repeated for the two functional runs (REST1, REST2) such that an individual had two functional connectivity matrices which reflect their connectivity profiles. We eliminated some edges in the functional connectivity matrices for each participant due to the lack of spatial coverage (i.e., zero values) across the whole brain. We also removed duplicate edges by considering only the upper triangular part of the functional connectivity matrices. There were 25,200 distinct edges that remained in REST1 and REST2.

### 2.7 Functional Connectivity: Scrubbing

To identify motion-corrupted time points in the fMRI time series, a volume censoring strategy commonly referred to as “scrubbing” was applied (**Figure 1**; Power et al., 2015; Power et al., 2014; Power et al., 2013). For each participant, a contaminated time point *T* is identified in their fMRI time series if its rmsFD > 0.20 mm. Once *T* has been identified, the preceding time point (*T* – 1) and two succeeding time points (*T* + 1, *T* + 2) are censored in the fMRI time series. By removing a total of four time points for each *T* in the fMRI time series, the residual motion that persists in the neighboring time points of *T* is minimized.

**Figure 1.**
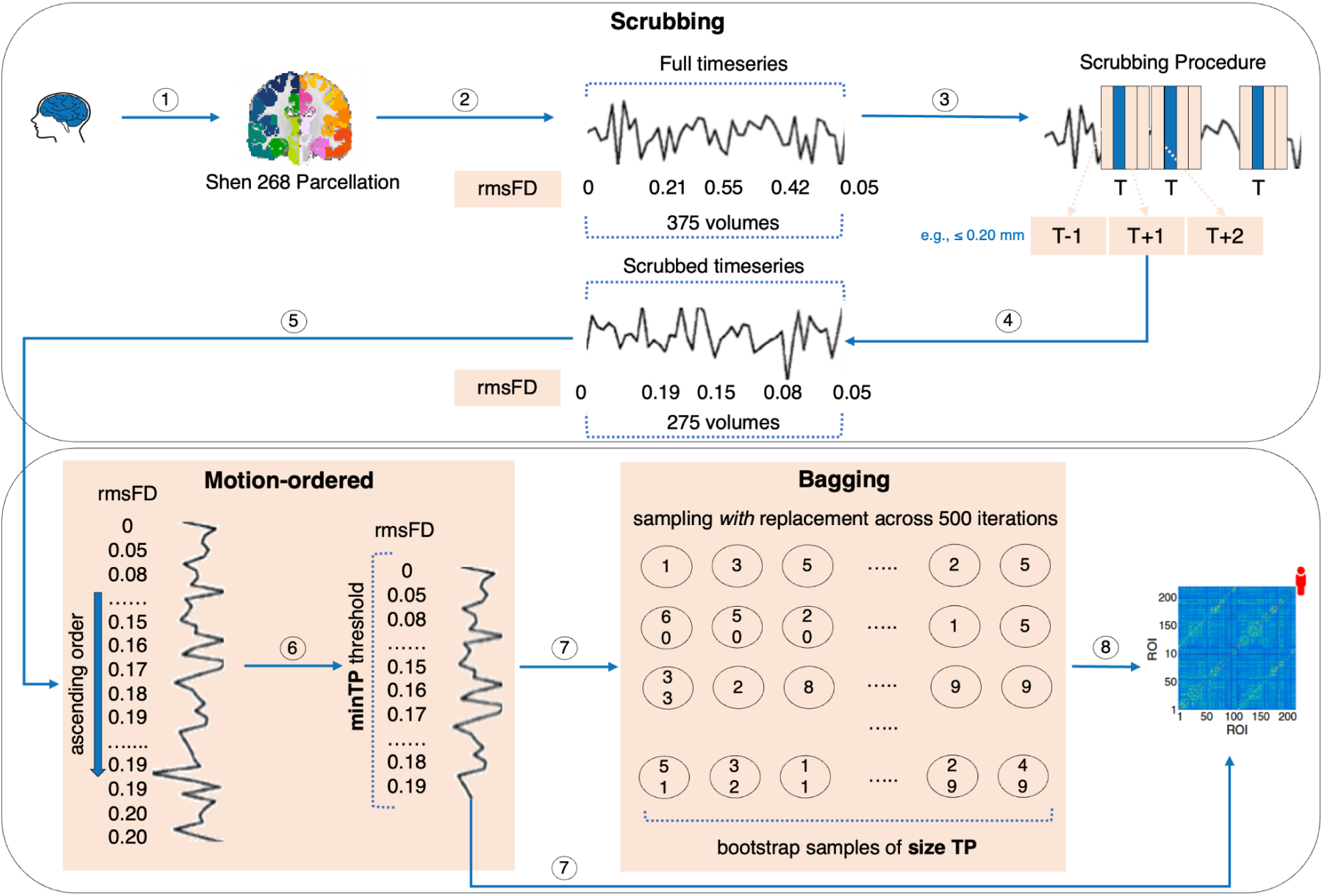
Schematic describing the motion-ordered and bagging procedures. The Shen 268 parcellation scheme is applied to extract the fMRI time series of all the participants. When scrubbing is performed, a motion-corrupted time point (*T*) is identified in the fMRI time series if its rmsFD > 0.20 mm. After censoring that time point, its preceding (*T* – 1) and two succeeding (*T* + 1, *T* + 2) time points are also removed. Time points are then ordered according to their rmsFD values, and the top minTP least motion-contaminated time points are used to compute functional connectivity (motion-ordered functional connectivity matrix). For each participant, the functional connectivity matrix is computed using the motion-ordered time series. Bagging is performed using the scrubbed time series by selecting the least motion-corrupted time points (ordered by their rmsFD values) that match a predefined threshold denoted by minTP, and bootstrapping samples, of a given size TP, *with* replacement, from the motion-limited time points across 500 iterations, and computing functional connectivity. For each participant, the mean bagged functional connectivity matrix is computed by averaging the resultant 500 functional connectivity matrices (bagged functional connectivity matrix).

### 2.8 Functional Connectivity: Motion-ordering and Bagging Approaches

Using the scrubbed time series, we employed the motion-ordered and bagging approaches depicted in **Figure 1** respectively to select a subset of the least motion-corrupted time points of length **minTP** and bootstrapped samples of length **TP** time points from the subset of least motion-corrupted time points, denoted by **minTP**. First, for each participant, we reordered their scrubbed time series according to rmsFD (i.e., following the method described by Fair et al., 2020), and retained **minTP** time points exhibiting the least motion — a procedure we refer to as motion-ordered approach. Applying the **minTP** threshold here is an important step that prevents varying numbers of time points being retained across participants, which could introduce bias. For each participant, a functional connectivity matrix was obtained by computing the pairwise correlation between the motion-ordered time series.

Next, we performed bagging by selecting 500 bootstrapped samples of length **TP** *with* replacement from the minTP-matched time points, and a functional connectivity matrix indexing the pairwise correlation between all time series was computed. For each participant, a mean bagged functional connectivity matrix was then obtained by averaging the functional connectivity matrices across the 500 samples. We further evaluated the importance of bagging by comparing results to those obtained using only a single sample (iteration) of length TP (relative to 500) — a procedure we refer to as “one-sample bootstrap”.

The approach to bagging taken here is more similar to that of Biswal et al. (2001), who subsampled task-based fMRI time series *without* replacement, than that of Bellec et al. (2010) and Nikolaidis et al. (2020), who used the circular block bootstrap. The circular block bootstrap is considered preferable because it better allows for temporal autocorrelation in the time series, which is known to contribute to functional connectivity (Arbabshirani et al., 2014) and fingerprint ID accuracy (Shinn et al., 2023). Here, due to the prioritization of low-motion time points, and the exclusion of the preceding and two succeeding time points after a motion-contaminated time point, it was not possible to apply circular block bootstrap while still retaining sufficient time points for the calculation of functional connectivity. Since all preprocessing, including confound regression and temporal filtering, was performed in the usual manner, we may expect only minimal impact of disrupted temporal autocorrelation, although the literature is mixed on whether this would be a positive (e.g., Arbabshirani et al., 2014) or negative (e.g., Shinn et al., 2023) outcome. Functional connectivity as measured by correlation itself is invariant to order, as long as all time series are identically reordered.

### 2.9 Functional Connectome Fingerprinting

We next tested the impact of the motion-ordered and bagging approach on our ability to identify an individual based on their functional connectivity profiles, as previously described by Finn et al. (2015). The identification procedure was performed by creating a “database” **D** which stored the motion-ordered or bagged functional connectivity matrices of each individual from REST1. Iteratively, the motion-ordered or bagged functional connectivity matrix from a given individual from REST2 was selected, and this functional connectivity matrix was treated as the “target matrix” **T**. The target matrix **T** was then compared with each of the functional connectivity matrices in **D** to find the corresponding matrix that shared the highest Pearson correlation coefficient (*r*). Participant identification (ID) was computed using binary identification (BID) such that the ID was assigned a score of 1 if the predicted identity matched the true identity of the individual, otherwise the ID was given a score of 0. The BID accuracy was computed as the percentage of individuals who were successfully identified from *N*. A mean BID accuracy was obtained by exchanging the **D**↔**T** configurations. The entire process was repeated for the standard and one-sample bootstrap approaches. Mean BID accuracies were then plotted as a function of time points ranging from 60 to the full time series length.

### 2.10 Brain-behavior Relationships

#### Standard Approach

To first establish a brain-behavior relationship to use in further validation analyses, we computed partial Spearman’s Rank correlations (r_s_) between functional connectivity and age at the edge level using the full time series for participants in the main sample of *N* = 255 who exhibited rmsFD < 0.20 mm, while treating sex and head motion (indexed by rmsFD) as covariates. The edge that shared the strongest brain-behavior relationship (highest r_s_) was selected after applying a False Discovery Rate (FDR) to correct for multiple comparisons (across 25,200 edges). Following the logic of Marek et al. (2022), we then established the robustness of this brain-behavior relationship across different sample sizes. We randomly selected samples of the *N* = 255 low-motion participants at 13 intervals: *N* ɛ {20, 40, 60, 80, 100, 120, 140, 160, 180, 200, 220, 240, 255}, with each sample being bootstrapped *without* replacement over 500 iterations. For each sample size, the mean brain-behavior r_s_ and 95% CIs were plotted as a function of *N*. We refer to this procedure as the *standard* approach.

#### Motion-ordered and Bagging Approaches

We repeated the brain-behavior analyses for the same edge by calculating r_s_ between functional connectivity and age using the motion-ordered and bagging approach applied to the same *N* = 255 participants, while treating sex and head motion (indexed by rmsFD) as covariates, that shared the highest correlation strength with the standard approach. The robustness of the brain-behavior relationship for the motion-ordered and bagged functional connectivity was computed using the same bootstrapping approach and 13 sample sizes described above, and the mean brain-behavior r_s_ and 95% CIs were plotted as a function of *N*. We compared the performance of the standard, motion-ordered, and bagging approaches by computing the areas under the curve (AUC) for the 95% CIs, such that a lower AUC indicates tighter 95% CIs. To compare the difference in AUC between the three approaches, we computed the difference in AUC, as denoted by ΔAUC. We also examined the ΔAUC to compare the performance of bagging and one-sample bootstrap.

### 2.11 Reducing exclusion of high-motion participants from connectome-based fingerprinting and brain-behavior analyses

To assess whether our motion-ordered and bagging methods can enable the rescue or retention of data from participants who would otherwise be excluded for exceeding motion thresholds, we computed fingerprint ID accuracies and brain-behavior r_s_ using the motion-ordered and bagged functional connectivity matrices derived from the scrubbed time series of all participants (*N* = 423; i.e., we did not apply the initial motion threshold of rmsFD < 0.20 mm). We first performed connectome-based fingerprinting as described in Section 2.9 using the motion-ordered and bagged functional connectivity matrices from REST1 and REST2, and including all participants from the *N* = 423 sample who had at least **minTP** uncorrupted time points available. The mean BID accuracies were plotted as a function of time point (TP) ranging from 60 to the full time series length. Subsequently, brain-behavior analyses described in Section 2.10 were repeated for a sample that includes these high-motion participants who were originally excluded from our analyses. We also tested for differences between these “rescued” high-motion participants and the group of “low-motion” participants originally included in analyses, based on several demographic characteristics (e.g., age, sex).

Finally, we performed two further assessments of reproducibility of our motion-ordered and bagging approaches. First, we assessed whether the motion-ordered and bagging approaches would enable the retention of high-motion participants who would have otherwise been discarded from brain-behavior analyses according to the application of a particularly stringent motion threshold of rmsFD < 0.08 mm, as previously employed by Marek et al. (2022). Second, we investigated the impact of the motion-ordered and bagging approaches on brain-behavior associations for the top 10 edges, ordered by their correlation strengths (r_s_) after applying an FDR threshold to correct for multiple comparisons (across 25,200 edges). Given that the edges that share a behavioral relevance with age may differ in their magnitude and direction, we assessed whether the two approaches can produce robust brain-behavior relationships regardless of the direction of specific edges in the functional connectome. We computed the AUC for the motion-ordered and bagging approaches across the different edges in the functional connectome. The mean brain-behavior r_s_ and 95% CIs were also plotted as a function of *N*.

To promote further assessment and application of our approach, all Python code used in this study is available in our GitHub repository (https://github.com/JRam02/bagging); further details are provided in the Data and Code Availability statement.

## 3 Results

### 3.1 Applying motion-ordered and bagging approaches for connectome-based fingerprinting

To test whether the motion-ordered and bagging approaches generate reproducible functional connectivity matrices, we computed fingerprint ID accuracies for the HBN dataset, after applying a rmsFD < 0.20 mm threshold to exclude participants with excess motion. Given that fingerprint ID accuracy is usually computed with the full time series, and tends to increase with longer time series (Vanderwal et al., 2021; Venkatesh et al., 2020; Vanderwal et al., 2017; Finn et al., 2015), we expectedly found that the BID accuracies were optimal (77-79%) with the standard approach (i.e., the scrubbed, full time series; **Figure 2A-B**). In the first scenario, the low-motion participants had at least 200 least motion-limited time points and the sample size was reduced to 239. When we applied the motion-ordered approach, BID accuracies were good (71-79%) with varying time points ranging from 60 to 200 (**Figure 2A**). Similarly, when we took the same subsets of motion-ordered time points but additionally applied bagging by varying the number of bootstrap time points from 60 to 200, fingerprint accuracies remained good (77-78%; **Figure 2A**). Importantly, the motion-ordered and bagging approaches produced similar fingerprint ID accuracies relative to the standard approach (375 time points). In the second scenario, we lowered the participants’ least motion-limited time points to 60 and the sample size increased to 255. To prevent a change in sample size, the motion-ordered approach was performed by using only the 60 motion-limited time points and BID accuracy remained good (66%; **Figure 2B**). Bagging also generated good BID accuracies (64-67%) with varying time points ranging from 60 to 200. In this scenario, time points above 60 were oversampled (**Figure 2B**). Although there was a decrease in accuracy overall, which may be driven by the shorter time series, using as few as 60 of the total 375 time points largely preserved BID accuracy and enabled identification of individuals based on their functional connectomes. There was also a clear benefit of computing the functional connectome on the basis of bootstrapped data in the two scenarios — when BID accuracies were calculated for functional connectomes computed using only one sample (rather than 500) was taken, they were lower across different number of bagged time points (**Figures 2A-B**).

**Figure 2.**
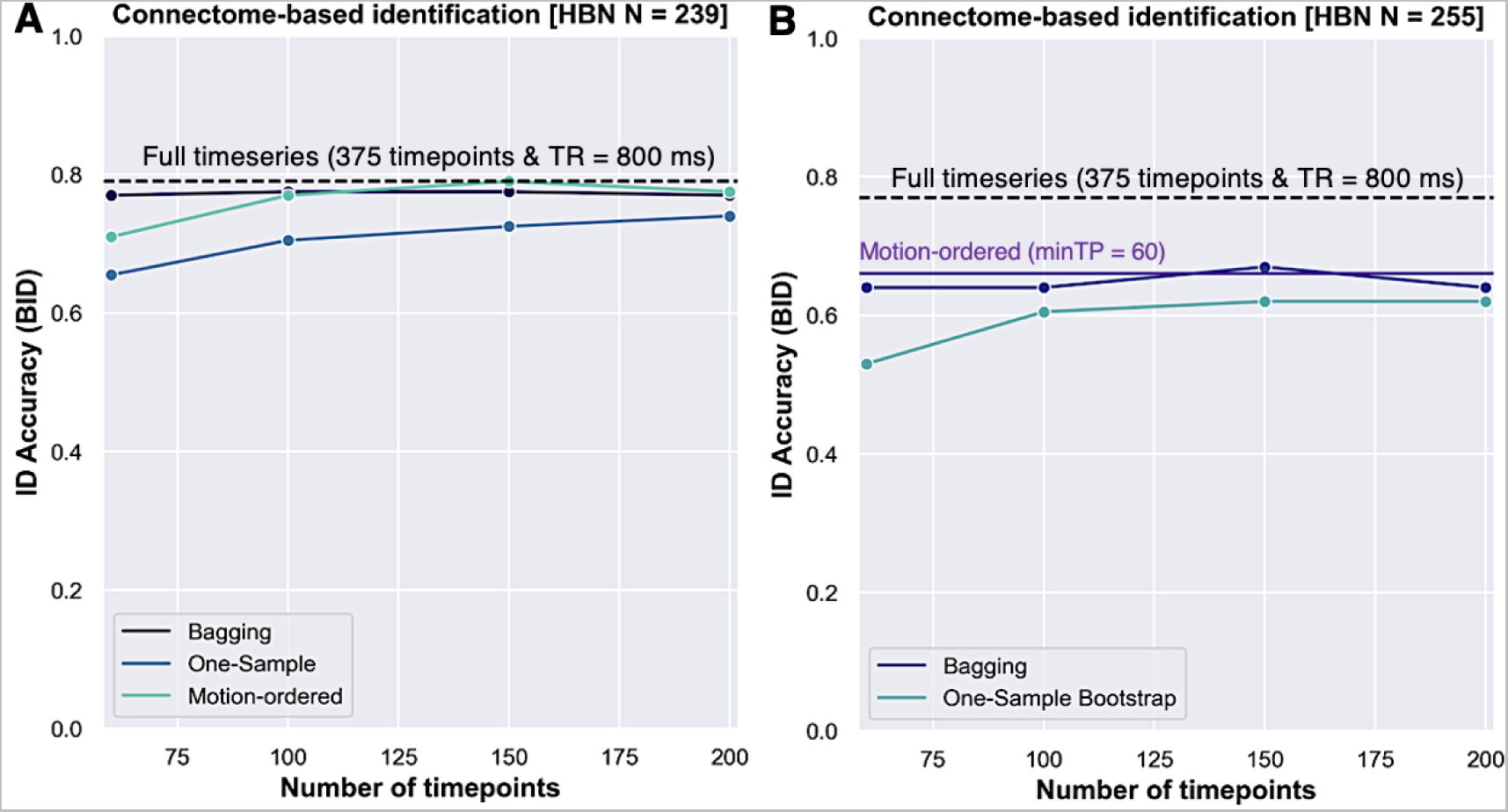
Applying motion-ordered, bagging, and one-sample bootstrap approaches to perform connectome-based fingerprint identification. Fingerprint accuracy measured by BID was performed by excluding “high-motion” participants with a rmsFD > 0.20 mm. (A) For the motion-ordered approach, BID accuracy was computed using the motion-limited time series by varying minTP from 60 to 200. For bagging and one-sample bootstrap, the BID accuracies were computed using the motion-limited time series with minTP = 200 as a function of time points ranging from 60 to 200. In this scenario, there was no oversampling of bagged time points. After applying a rmsFD < 0.20 mm to exclude “high-motion” participants in addition to setting minTP = 200, the sample size was reduced to 239. (B) For the motion-ordered approach, the BID accuracy was computed using the motion-limited time series with minTP = 60. For bagging and one-sample bootstrap, the BID accuracies were computed using the motion-limited time series with minTP = 60 as a function of time points ranging from 60 to 200. In this scenario, there was oversampling of bagged time points. After applying a rmsFD < 0.20 mm to exclude “high-motion” participants in addition to setting minTP = 60, the sample size increased to 255. Solid black dotted line represents the BID accuracy obtained from the standard approach using the full time series corresponding to 375 timepoints with TR = 800 ms (5 min).

### 3.2 Standard brain-behavior relationships as a function of sample size

We computed the brain-behavior associations by performing partial Spearman’s Rank correlation (r_s_) between functional connectivity and age while treating head motion (indexed by rmsFD) and sex as covariates. We first employed the standard approach, which uses the full time series from REST1 (*N* = 255). The edge exhibiting the strongest brain-behavior relationship (highest r_s_) between functional connectivity and age was selected for this analysis. Following Marek et al. (2022), plotting the brain-behavior associations between functional connectivity and age as a function of sample size (*N*), revealed an expected tightening of the 95% CIs as the sample size increased from small (*N* = 20) to large (*N* = 255) (**Figure 3**). 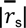 was 0.35 across *N*s.

**Figure 3.**
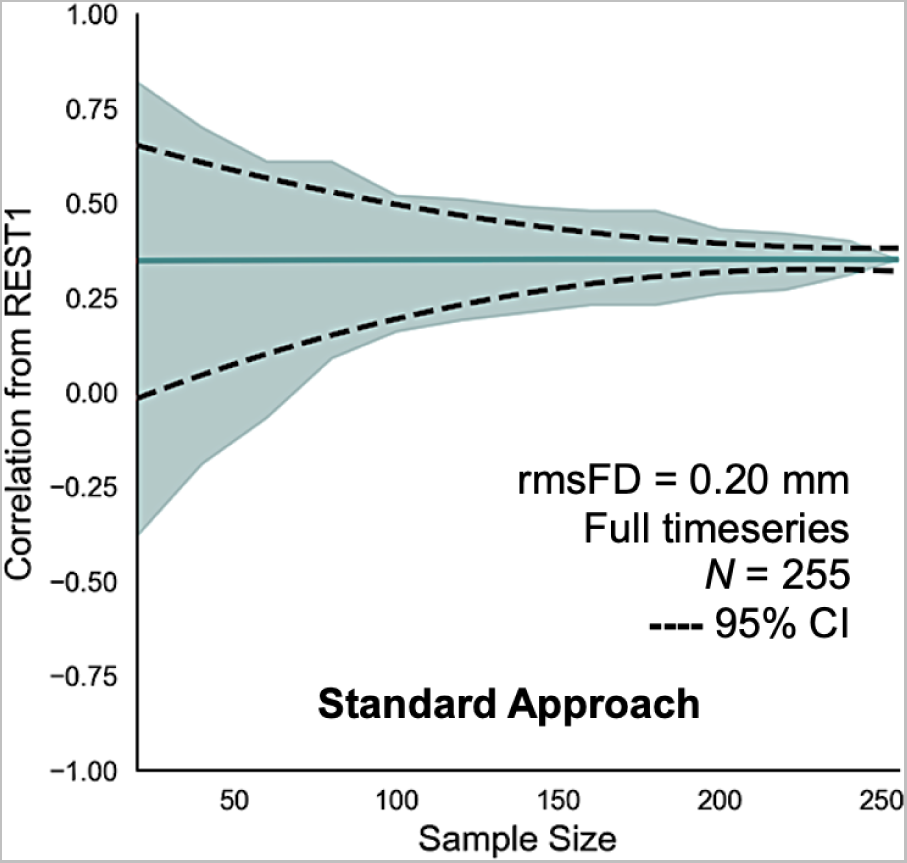
Correlations between functional connectivity and age obtained from REST1 using the standard approach as a function of sample size. Solid teal line shows the mean correlations derived from 500 bootstrap samples for a given sample size. Teal shading denotes the minimum and maximum correlations across 500 bootstrap samples for a given sample size. Black dotted line represents the lower and upper bounds of the 95% CIs. After applying a rmsFD < 0.20 mm to exclude “high-motion” participants, the sample size was reduced to 255.

### 3.3 Motion-ordered and bagged brain-behavior relationships as a function of sample size

We repeated the analysis of brain-behavior relationship strengths as a function of sample size using the bagging approach. The brain-behavior relationships between functional connectivity and age were computed using the same edge as in Section 3.2. In the first instance, we set minTP = 200, i.e., we selected the 200 least motion-corrupted time points from each participant’s scrubbed time series, based on their rmsFD values. As a result of this requirement, the sample size was reduced to *N* = 239. To enable fair comparisons of the performance of bagging with the standard approach, we repeated the brain-behavior analyses using the standard approach for *N* = 239. Using the standard approach, we observed similar tightening of the 95% CIs as the sample size increased from typical (*N* = 20) to large (*N* = 239), and 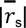 was 0.37 (**Figure 4A**). Both the motion-ordered (minTP = 200) and bagging (minTP = 200 and TP = 60) achieved robust correlations between functional connectivity and age which were comparable to those obtained using the standard approach (motion-ordered: 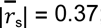 ; bagging: 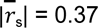 (**Figures 4B-C**).

**Figure 4.**
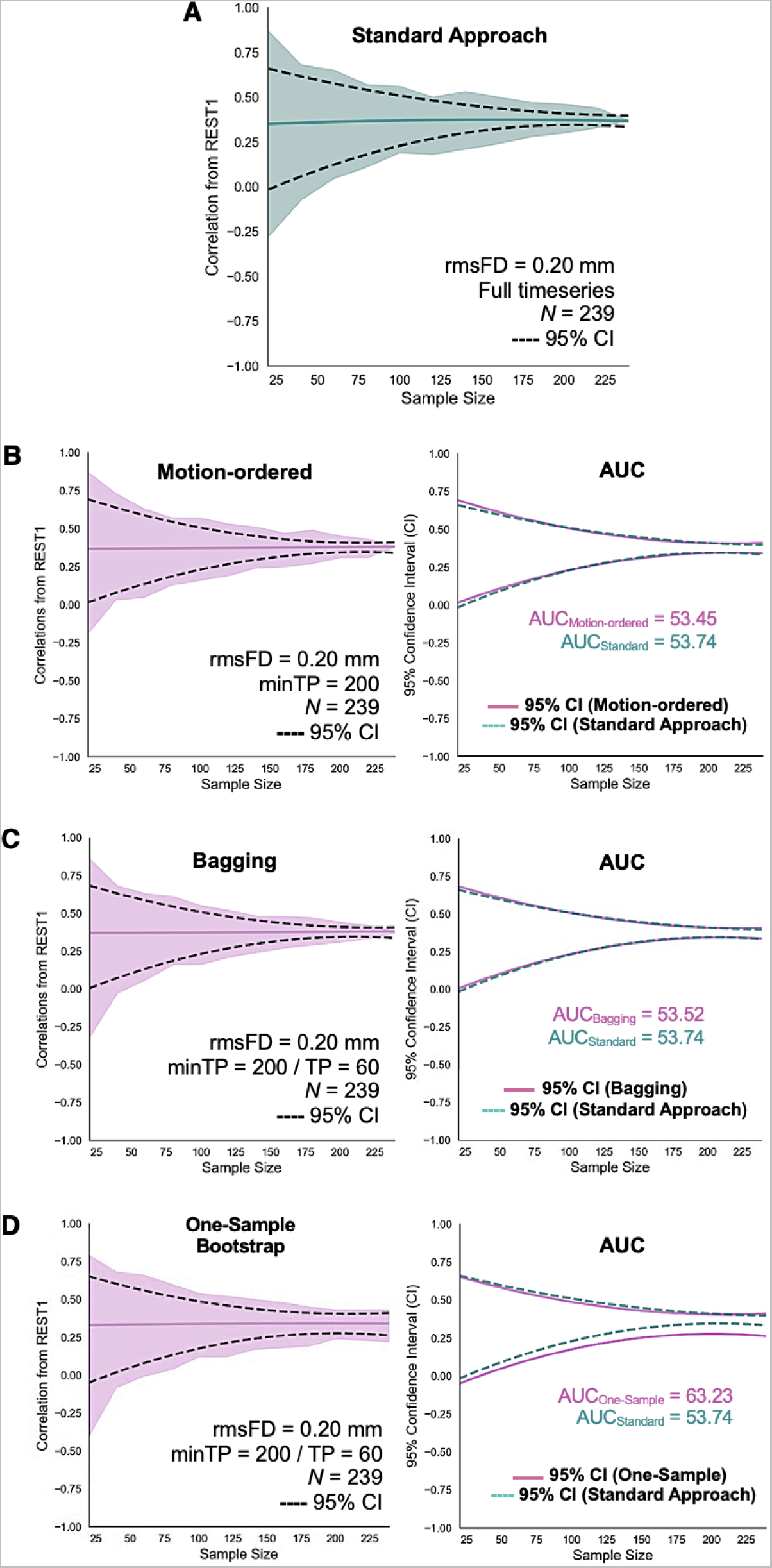
Correlations between functional connectivity and age obtained from REST1 as a function of sample size using the motion-ordered, bagging, and one-sample bootstrap approaches. (A) Correlations between functional connectivity and age obtained from REST1 using the standard approach as a function of sample size. The sample size was set to 239 to facilitate the comparison of the standard approach with the motion-ordered and bagging approaches. (B) Correlations between functional connectivity and age obtained from REST1 as a function of sample size using the motion-ordered approach during which participants have least amount of motion-corrupted time points that match the minTP threshold set at 200. (C) Correlations between functional connectivity and age obtained from REST1 as a function of sample size using bagging during which participants have least amount of motion-corrupted time points that match the minTP threshold set at 200 and number of bagged time points, i.e., TP set at 60. (D) Correlations between functional connectivity and age obtained from REST1 as a function of sample size using one-sample bootstrap during which participants have least amount of motion-corrupted time points that match the minTP threshold set at 200 and number of bagged time points, i.e., TP set at 60. The comparisons of the 95% CIs between the standard approach and motion-ordered, bagging, and one-sample bootstrap approaches are also shown with the area under curve (AUC). Dotted teal line represents the 95% CIs derived from the standard approach. Solid orchid lines represent the 95% CIs derived from the motion-ordered, bagging, and one-sample bootstrap approaches.

These observations were confirmed by the AUC analysis. The ΔAUC, which quantifies the magnitude of change (i.e., the difference) in AUC between the standard and motion-ordered approaches (ΔAUC = 0.54%) was small (**Figures 4A-B**). A similar difference was noted when comparing the AUCs between the standard and bagging approaches (ΔAUC_TP60_ = 0.40%) (**Figures 4A & C**). In contrast, Δ AUC between standard and one-sample bootstrap approaches was associated with significant widening of the 95% CIs (ΔAUC_TP60_ = 17.7%) (**Figures 4A & D**). Similar observations were noted when comparing the AUC from one-sample bootstrap to the motion-ordered (ΔAUC = 18.3%) and bagging (ΔAUC_TP60_ = 18.1%) approaches (**Figures 4B-D**). It is important to note that the motion-ordered approach involved computing 13 intervals x 500 bootstrap *N* samples = 6,500 correlations whereas bagging involved computing 13 intervals x 500 bootstrap *N* samples x 500 bootstrap time points = 3.25 million correlations.

### 3.4 Robust brain-behavior associations with minimally bootstrap time points

To investigate the impact of the number of time points available for analysis, we reduced minTP from 200 to 60 for the motion-ordered and bagging approaches. For bagging, we further set TP to 20 which increased the sample size to *N* = 255. Even with as few as 60 total time points for the motion-ordered approach and bagging with 20 time points bootstrap-sampled, we observed robust brain-behavior associations (**Figure 5**). The standard approach yielded comparable brain-behavior associations with both the motion-ordered (ΔAUC = 2.9%) and bagging (ΔAUC_TP20_ = 5.5%) approaches as demonstrated by their AUCs. Increased variability in the brain-behavior correlations was observed with one-sample bootstrap relative to the motion-ordered (ΔAUC = 24.2%) and bagging (ΔAUC_TP20_ = 21.1%) approaches (**Figure 5**). Similar variability was noted when comparing the brain-behavior correlations obtained from one-sample bootstrap with the standard approach (ΔAUC_TP20_ = 27.8%). We also showed the influence of minTP and TP on the brain-behavior relationships as a function of sample size in **Supplementary Figure 1**. Similarly, the motion-ordered approach computed 13 intervals x 500 bootstrap *N* samples = 6,500 correlations whereas bagging computed 13 intervals x 500 bootstrap *N* samples x 500 bootstrap time points = 3.25 million correlations.

**Figure 5.**
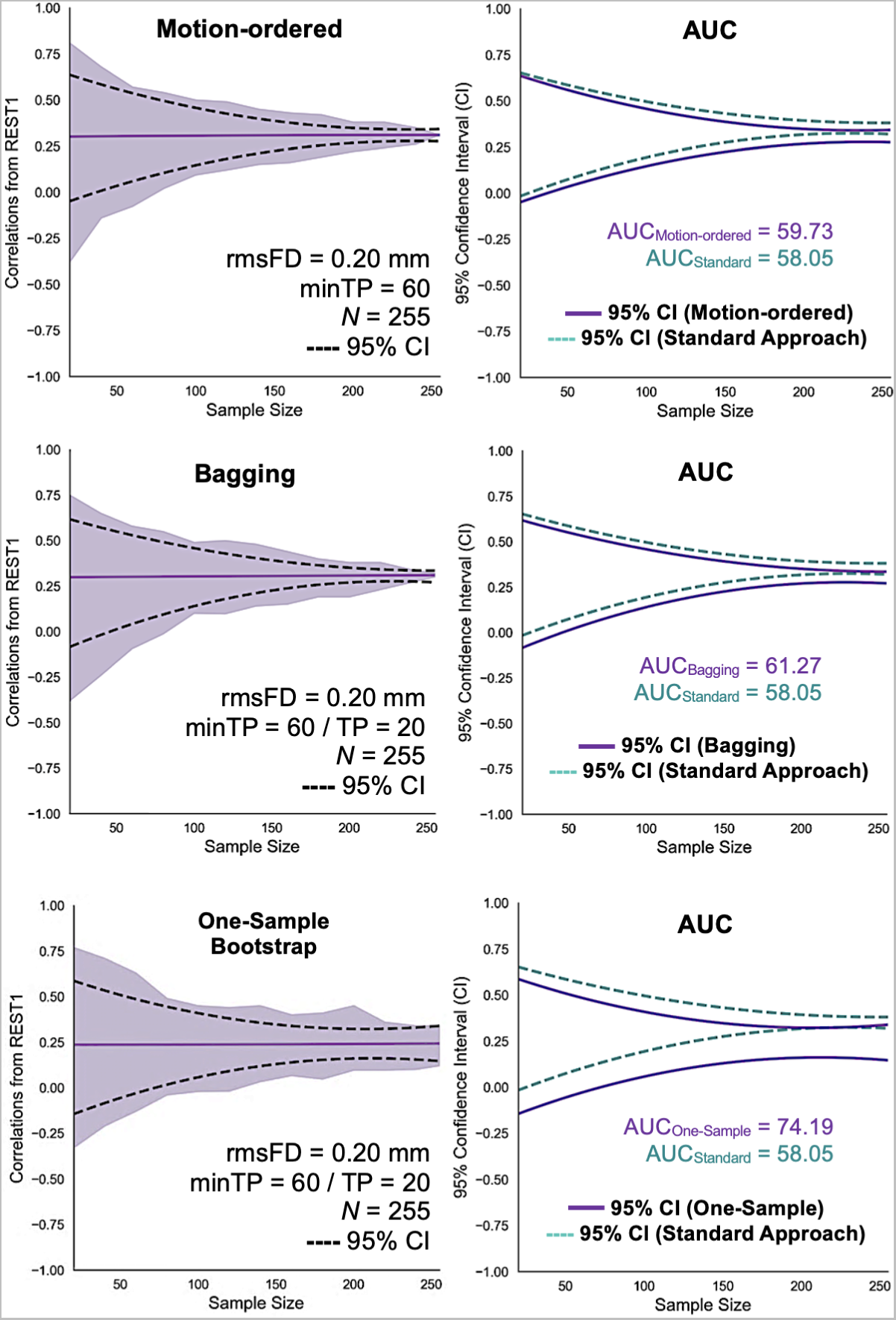
Correlations between functional connectivity and age obtained from REST1 as a function of sample size using the motion-ordered, bagging, and one-sample bootstrap approaches. For the three approaches, the motion-limited time series was set to a length of minTP = 60. For bagging and one-sample bootstrap, the number of bagged time points, i.e., TP was set to 20. That is, participants have minimally bootstrap time points that match the minTP threshold (e.g., 60) and number of bagged time points of length TP = 20. Solid purple line shows the mean correlations, purple shading denotes the minimum and maximum correlations, and black dotted line represents the lower and upper bounds of the 95% CIs. The AUC is displayed to compare the 95% CIs from the standard approach with the motion-ordered, bagging, and one-sample bootstrap approaches. Dotted teal line represents the 95% CIs derived from the standard approach. Solid purple lines represent the 95% CIs derived from the motion-ordered, bagging, and one-sample bootstrap approaches.

### 3.5 Applying motion-ordered and bagging approaches to perform connectome-based fingerprint identification with “high-motion” individuals

Until now, we employed the motion-ordered and bagging approaches to perform all the analyses after excluding participants with a rmsFD < 0.20 mm, resulting in the retention of ∼60% of the HBN (*N* = 255 of 423) participants. In this analysis, we examined whether the motion-ordered and bagging approaches would enable us to retain as many participants as possible without applying the initial rmsFD threshold. For the motion-ordered approach, we set minTP = 60, and for bagging, we set minTP = 60 and TP = 20, which helped retain ∼90% of the HBN (*N* = 379/423) participants. BID accuracies were optimal with the standard approach at 68% (**Figure 6**). However, the BID accuracies could be inflated due to the presence of motion in the full time series, especially from those who typically are labeled “high-motion”. When we applied the motion-ordered approach with minTP = 60, the BID accuracy reduced from 68% to 57% (**Figure 6**). Similarly, applying bagging with minTP = 60 and varying the number of bootstrap time points from 60 to 200, reduced BID accuracies from 68% to 58% (**Figure 6**). The one-sample bootstrap exhibited greater variability across different number of bagged timepoints (**Figure 6**).

**Figure 6.**
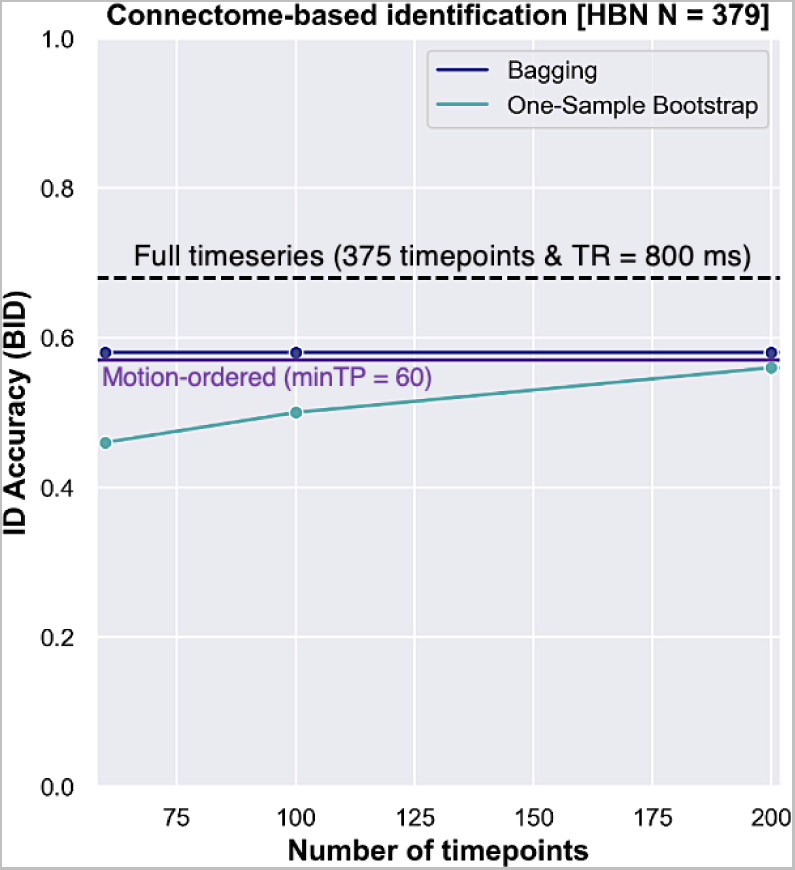
Applying motion-ordered, bagging, and one-sample bootstrap approaches to perform connectome-based fingerprint identification. For the motion-ordered approach, the BID accuracy was computed using the motion-limited time series with minTP = 60 by including the rescued “high-motion” participants. For bagging and one-sample bootstrap, the BID accuracies were computed using the motion-limited time series with minTP = 60 as a function of time points ranging from 60 to 200 by including the rescued “high-motion” participants. By including the “high-motion” participants on the basis of their 60 least motion-corrupted time points, the sample size increased to 379. Dotted purple line represents the BID accuracy obtained from the motion-ordered approach. Solid navy line denotes the BID accuracies obtained from bagging. Solid teal line corresponds to the BID accuracies obtained from one-sample bootstrap. Solid black dotted line represents the BID accuracy obtained from the standard approach using the full time series corresponding to 375 timepoints with TR = 800 ms (5 min).

### 3.6 Rescuing “high-motion” participants for brain-behavior associations

Including high-motion participants (*N* = 379) to compute the brain-behavior associations with the motion-ordered (minTP = 60) and bagging (minTP = 60 and TP = 20) approaches respectively yielded 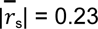 and 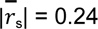 in addition to the expected narrowing of the 95% CIs with increasing sample sizes (**Figure 7**). We then compared the brain-behavior correlations from bagging and one-sample bootstrap with those obtained using the motion-ordered approach (**Figure 7**). The motion-ordered and bagging approaches produced similar narrowing of the 95% CIs with increasing sample sizes (ΔAUC = 0.96%), supporting the merits of sampling a subset of scrubbed time points (here, as little as 16% of the original time series) for high-motion participants. The one-sample bootstrap showed greater variability in the brain-behavior associations relative to the motion-ordered (ΔAUC = 17.2%) and bagging (ΔAUC_TP20_ = 18.3%) approaches. It is important to note that the motion-ordered approach computed 10 intervals x 500 bootstrap *N* samples = 5,000 correlations whereas bagging computed 10 intervals x 500 bootstrap *N* samples x 500 bootstrap time points = 2.5 million correlations.

**Figure 7.**
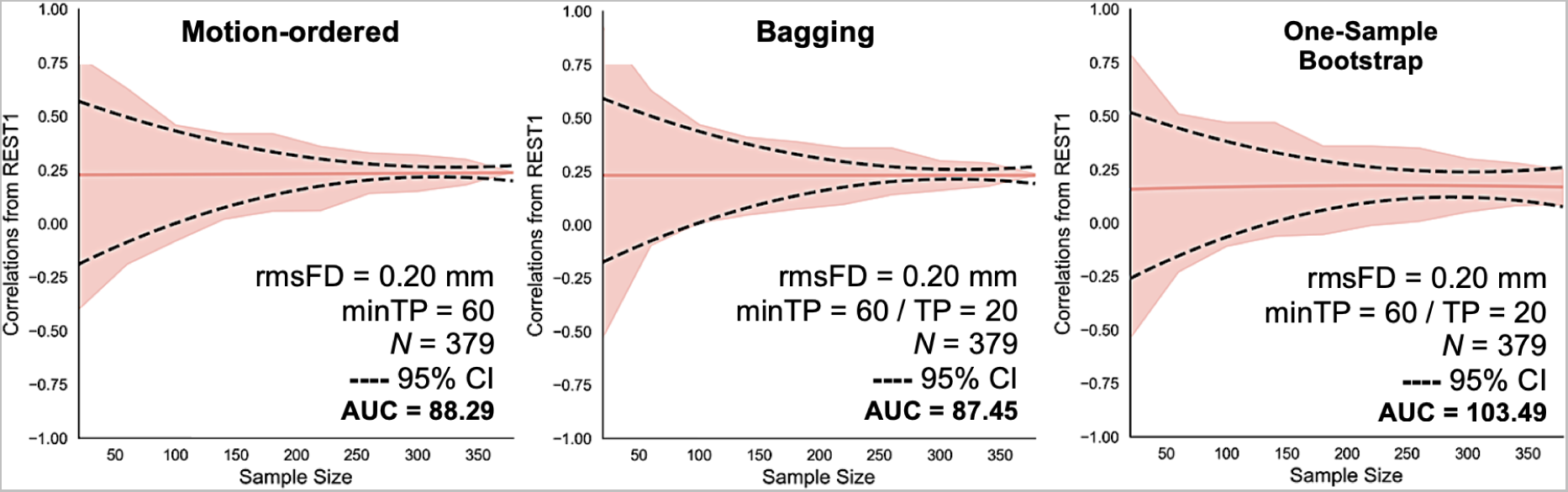
Formerly excluded “high-motion” participants can be included to compute brain-behavior associations with bagging. By applying the motion-ordered (minTP = 60), bagging (minTP = 60 and TP = 20), and one-sample bootstrap (minTP = 60 and TP = 20) approaches, the sample size increased to 379. The correlations between functional connectivity and age obtained from REST1 as a function of sample size are shown using the motion-ordered, bagging, and one-sample bootstrap approaches. The AUC is displayed for the motion-ordered, bagging, and one-sample bootstrap approaches. Solid pink line shows the mean correlations, pink shading denotes the minimum and maximum correlations, and black dotted line represents the lower and upper bounds of the 95% CIs.

Finally, we examined whether the “rescued” high-motion participants and the group of “low-motion” participants originally included in analyses, differed on several demographic characteristics (e.g., age, gender). Expectedly, relative to the low-motion participants, high-motion participants exhibited significantly greater average motion [rmsFD] for REST1 (Mann-Whitney U test, U = 1,870.5, *P <* 0.001) and REST2 (Mann-Whitney U test, U = 450.5, *P <* 0.001). Relative to the low-motion participants, high-motion participants were younger (mean age = 9.8 years vs 11.8 years; Mann-Whitney U test, U = 21,878.5, *P <* 0.001) and were twice as likely to be male (two-way Chi-Square test, 𝓧^2^ = 6.93, *P* = 0.008; odds ratio of male to female participants is 1.9). The two groups did not differ on the measures of clinical symptomatology we assessed (Child Behavior Checklist [CBCL] externalizing and internalizing; Mann-Whitney U test, *P* > 0.05), however (see **Supplementary Figures 2-3**).

### 3.7 Reproducibility of motion-ordered and bagging approaches across head motion thresholds and edges in the functional connectome

Up to this point, a head motion threshold of rmsFD < 0.20 mm was applied. Next, we evaluated the impact of a much stricter motion threshold, i.e., rmsFD < 0.08 mm, as recently utilized by Marek et al. (2022) to identify high-motion participants. First, we applied this threshold prior to computing the correlations with the motion-ordered (minTP = 60) and bagging (minTP = 60 and TP = 20) approaches, which resulted in the retention of only ∼18% of the sample (*N* = 77/423). Both the motion-ordered and bagging approaches produced similar brain-behavior relationships (motion-ordered: 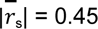 ; bagging: 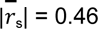) in addition to observing similar tightening of the 95% CIs as before (**Figure 8A**). Greater variability in the brain-behavior associations was observed when one-sample bootstrap was performed relative to the motion-ordered (ΔAUC = 68.6%) and bagging (Δ AUC_TP20_ = 69.8%) approaches (**Figure 8A**). Second, we retained all the participants with at least minTP = 60, which resulted in ∼22% of the sample size (*N* = 92/423) for the brain-behavior analyses. Similar brain-behavior relationships were observed for both the motion-ordered and bagging approaches (motion-ordered: 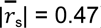; bagging: 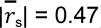) (**Figure 8B**). However, it is important to note that this strict rmsFD threshold retains fewer participants which can *inflate* the effect sizes of brain-behavior relationships due to smaller *N*s.

**Figure 8.**
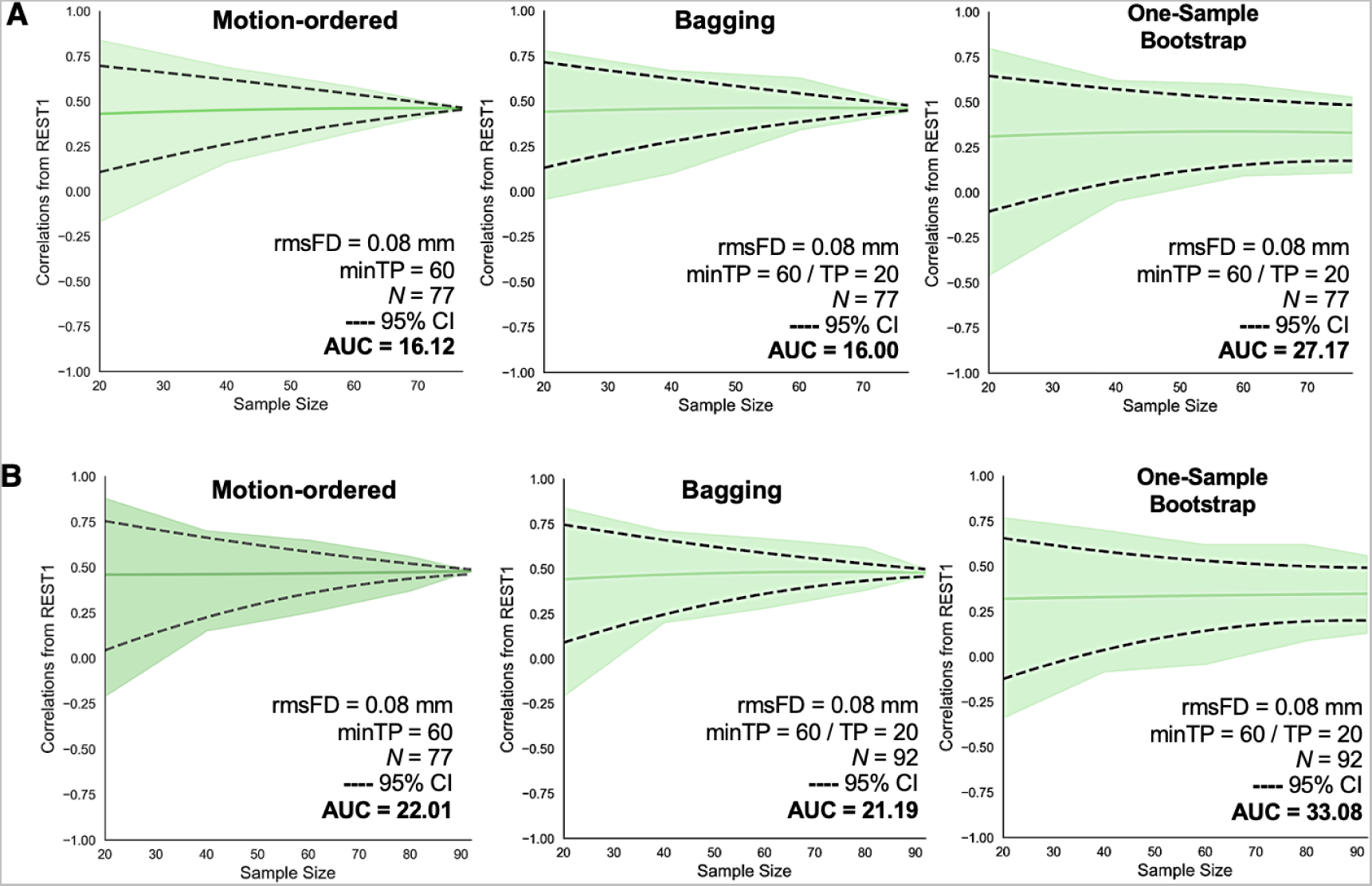
Impact of applying a strict head motion threshold on brain-behavior associations using the motion-ordered (minTP = 60), bagging (minTP = 60 and TP = 20), and one-sample bootstrap (minTP = 60 and TP = 20) approaches. (A) After applying a rmsFD < 0.08 mm to exclude “high-motion” participants, the sample size was reduced to 77. Correlations between functional connectivity and age obtained from REST1 as a function of sample size are shown using the motion-ordered, bagging, and one-sample bootstrap approaches. (B) After scrubbing all the participants’ time series with a rmsFD < 0.08 mm, the sample size was increased to 92. Correlations between functional connectivity and age obtained from REST1 as a function of sample size are shown using the motion-ordered, bagging, and one-sample bootstrap approaches. The AUC is displayed for the motion-ordered, bagging, and one-sample bootstrap approaches. Solid green line shows the mean correlations, green shading denotes the minimum and maximum correlations, and black dotted line represents the lower and upper bounds of the 95% CIs.

So far, we computed r_s_ using the standard, motion-ordered, and bagging approaches derived from the edge that shares the strongest correlation between functional connectivity and age. To test the reproducibility of the motion-ordered (minTP = 60) and bagging (minTP = 60 and TP = 20) approaches across different edges in the whole-brain functional connectome, we selected an additional 9 edges to compute the brain-behavior associations as a function of *N*. Both the motion-ordered and bagging approaches applied to data that included the previously excluded high-motion participants (*N* = 379) yielded reproducible brain-behavior relationships with AUCs (**Figures 9-10**). For the motion-ordered approach, the AUCs ranged from 81.81 to 87.12 (mean AUC ±SD = 84.61±2.20) and for bagging, the AUCs ranged from 80.72 to 85.53 (mean AUC±SD = 83.54 ±1.82).

**Figure 9.**
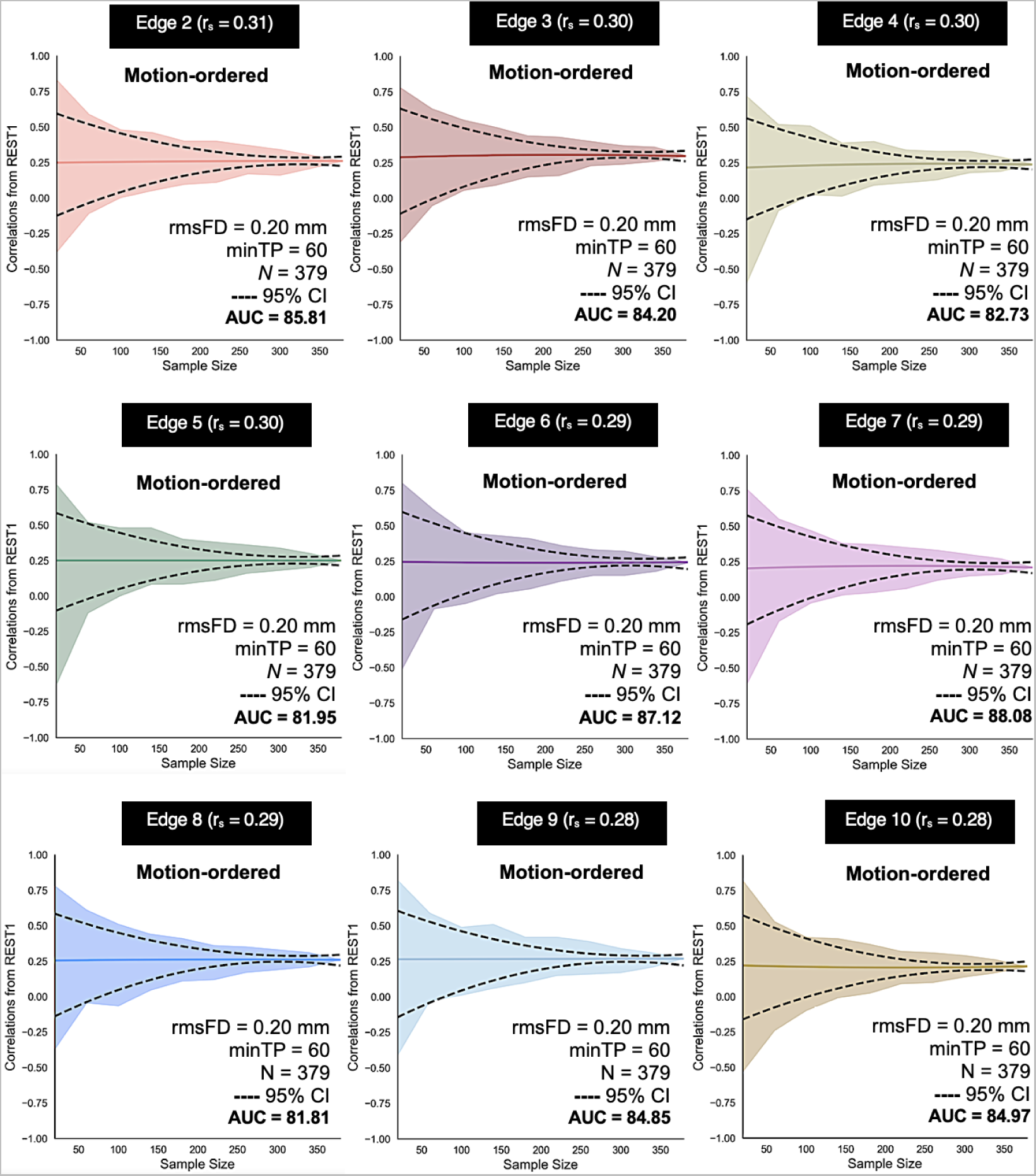
Reproducibility of motion-ordered approach (minTP = 60) including the “rescued” “high-motion” participants in deriving the associations between functional connectivity and age from an additional 9 edges in the whole-brain functional connectome. Correlations between functional connectivity and age obtained from REST1 as a function of sample size are shown using bagging across the 9 edges. The AUCs are displayed for bagging across the 9 edges. Solid colored lines show the mean correlations, colored shadings denote the minimum and maximum correlations, and black dotted lines represent the lower and upper bounds of the 95% CIs. After retaining the “high-motion” participants, the sample size increased to 379.

**Figure 10.**
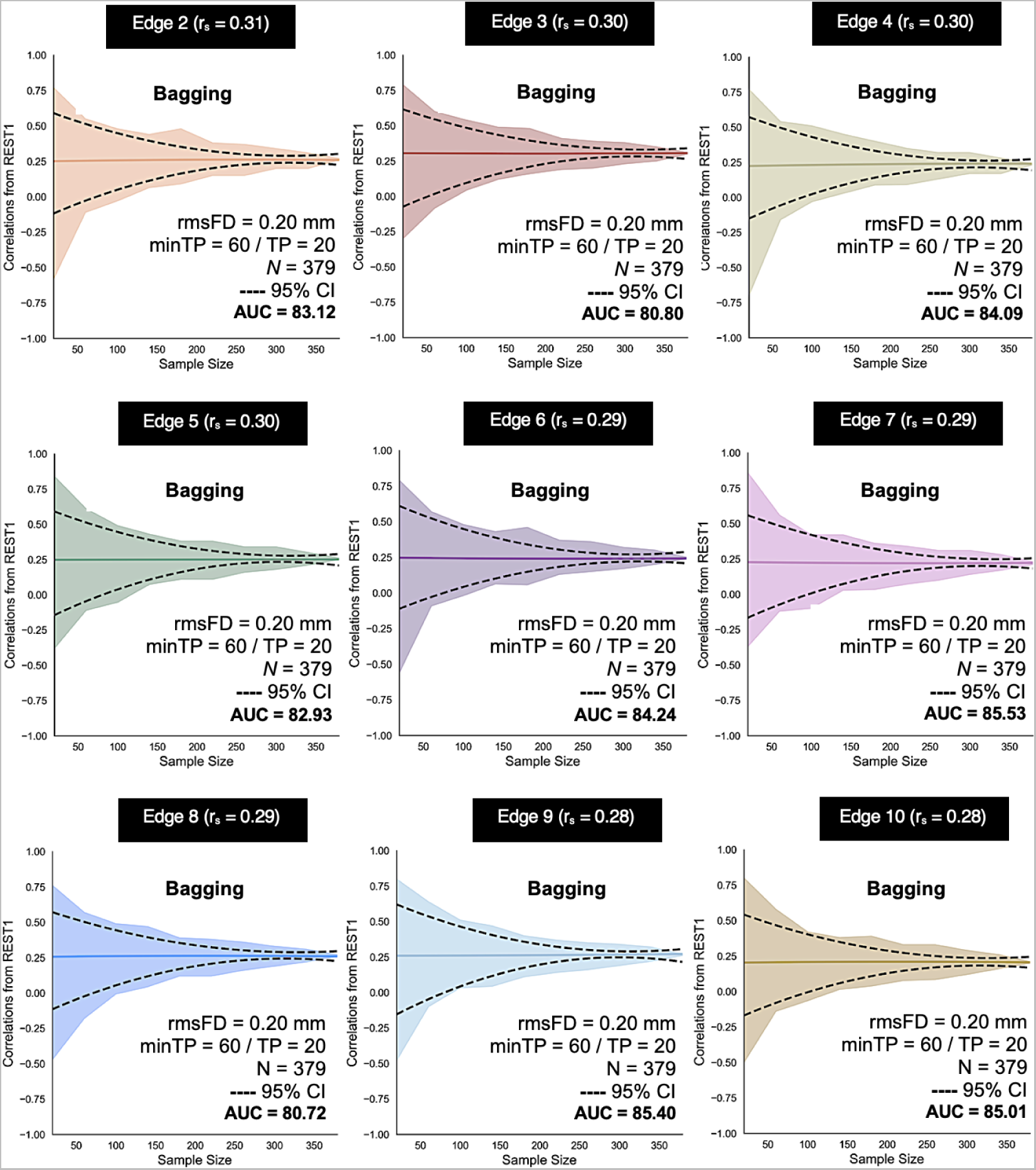
Reproducibility of bagging (minTP = 60 and TP = 20) including the “rescued” “high-motion” participants in deriving the associations between functional connectivity and age from an additional 9 edges in the whole-brain functional connectome. Correlations between functional connectivity and age obtained from REST1 as a function of sample size are shown using bagging across the 9 edges. The AUCs are displayed for bagging across the 9 edges. Solid colored lines show the mean correlations, colored shadings denote the minimum and maximum correlations, and black dotted lines represent the lower and upper bounds of the 95% CIs. After retaining the “high-motion” participants, the sample size increased to 379.

### 3.8 Adjudicating between approaches

Given that the motion-ordered and bagging approaches exhibited equivalent performance in terms of both fingerprint ID accuracy, the stability of brain-behavior relationships across different sample sizes and edges in the functional connectome, and the rescue of high-motion participants, we suggest that computing functional connectivity on the basis of timeseries that have been subsetted to include the lowest-motion time points (i.e., the *motion ordered* approach) is preferable, due to its lower computational demands.

## 4 Discussion

To achieve robust and reproducible brain-behavior relationships in neuroimaging-based studies, we need large sample sizes that can provide adequate statistical power while applying rigorous head motion thresholds that guarantee data quality. These imperatives can come into conflict: applying strict head motion thresholds reduces sample size, sometimes substantially, particularly in developmental or clinical cohorts. To address this conflict, the present study assessed the impact of two head motion mitigation techniques, a motion-ordering and a motion-ordering+resampling (i.e., bagging), on functional connectome fingerprinting accuracy and brain-behavior associations in a sample of developing youth. First, we demonstrated that the motion-ordered and bagging approaches produce reproducible functional connectivity matrices that successfully differentiate an individual from a group — *with* and *without* the inclusion of the high-motion individuals’ bootstrap motion-limited time points in a developmental sample. Second, we showed that functional connectomes computed using both the motion-ordered and bagging approaches retain validity, yielding robust brain-behavior correlations that vary as a function of sample size, even when computed using as few as 60 time points (i.e., 16% of the original fMRI time series). Third, we showed that the motion-ordered and bagging approaches improve inclusivity by enabling the “rescue” of data from high-motion participants — analyses including these individuals yielded brain-behavior association patterns that were comparable to analyses that excluded them. Last, we demonstrated that the motion-ordered and bagging approaches generate reproducible brain-behavior associations across different head motion thresholds and edges in the whole-brain functional connectome. Given equivalent performance of the two approaches, we recommend the motion-ordered approach to enable data retention and to enhance inclusivity for functional connectivity analyses.

### Utility of motion-ordered and bagging approaches

In our first assessment of the utility of the motion-ordered and bagging approaches, we focused on reproducibility. We examined the impact of the two approaches on connectome-based identification in our developmental sample. We demonstrated that motion-ordered and bagged functional connectomes can successfully identify developing participants using as few as 60 time points, which represents scan lengths < 1 min. There was an overall decrement in ID accuracy, which is to be expected, given that previous studies have shown that the ability to identify someone based on their functional connectivity fingerprints increases in association with scan duration (Vanderwal et al., 2021; Venkatesh et al., 2020; Vanderwal et al., 2017; Finn et al., 2015). Nonetheless, here, using even minimal time series (minTP = 60), fingerprint ID accuracy was largely preserved both *with* and *without* the high-motion participants. While larger *N*s typically decrease fingerprint ID accuracy (Li et al., 2021; Waller et al., 2017), this should be weighed against the possible advantages of participant inclusivity. Indeed, we observed that the high-motion participants were younger than the low-motion participants, which is in line with previous work examining the characteristics of participants who are routinely excluded from fMRI-based studies due to excess motion (Frew et al., 2022; Vanderwal et al., 2019; Vanderwal et al., 2015). There is considerable scope for further analyses to probe the opportunities and limitations of the motion-ordered and bagging approaches, but we nevertheless suggest that they may be applicable as a reproducible technique to increase *N* and minimize motion-related data loss, particularly in challenging developing populations for which the connectome distinctiveness is low (Dufford et al., 2021).

### Validity of motion-ordered and bagging approaches

Our second assessment of the motion-ordered and bagging approaches focused on validity — we examined the impact of the motion-ordered and bagging approaches on estimates and CIs for brain-behavior relationships as a function of sample size. When the two approaches were applied *with* and *without* the inclusion of high-motion participants, increasingly robust brain-behavior associations were observed as the sample size was increased from small to large. This is to be expected due to sampling variability; using two metrics, the CI and AUC, we showed that, as observed for the standard, scrubbed but unbagged data, variability decreased as the number of participants being sampled increased. This is fully in line with previous reports and demonstrates that brain-behavior correlations are inflated with typical (e.g., *N* = 20) compared to larger (e.g., *N* = 255) sample sizes because small samples are more prone to exhibit random variations in these associations, potentially overestimating effect sizes, and limiting reproducibility (Marek et al., 2022). Although the magnitude of the change in CIs between the standard, motion-ordered, and bagging approaches, as quantified by the difference in AUC, varied across edges, the two proposed techniques had a relatively small impact on the robustness of brain-behavior relationships, suggesting that it does not negatively impact validity. Our analyses also demonstrated that developing youth who exhibit relatively high levels of head motion — who would normally be excluded from analyses based on head motion thresholds — have usable data that are meaningful for brain-behavior associations. The inclusion of these participants, who tend to be younger and at risk of severe clinical symptoms, as our analyses and previous work (e.g., Vanderwal et al., 2019; Vanderwal et al., 2015) have shown, may increase both the validity and reproducibility of brain-behavior analyses by mitigating the range restriction that occurs when they are excluded. This presents a promising avenue for future research.

### Generalizability of motion-ordered and bagging approaches

We further showed that the motion-ordered and bagging approaches yielded reproducible brain-behavior associations across different edges in the functional connectome. This suggests that the benefits of the two approaches are not restricted to specific connections or relationships. Nonetheless, future work should evaluate their generalizability using other behavioral phenotypes. Age was chosen as the behavior of interest to perform brain-behavior associations as its relationship with functional connectivity is robust and has been well-studied in neurodevelopment (Grayson et al., 2017; Geerlings et al., 2015; Song et al., 2014; Di Martino et al., 2014a; Menon, 2013; Uddin et al., 2011; Park and Reuter-Lorenz, 2009; Fair et al., 2009; Fair et al., 2007). For example, the motion-ordered and bagging approaches could be applied to study the brain-behavior associations with measures that index cognitive ability and general psychopathology — which have recently been shown to generate reproducible brain-wide phenotypic associations though only in the context of very large sample sizes (Marek et al., 2022). Moreover, achieving generalizable brain-behavior associations in both small and large datasets remains important, particularly in developmental cohorts typically characterized by excess head motion, such as the ABCD Study® (Casey et al., 2018).

### Comparing motion-ordering and bagging

Given previous work demonstrating the utility of subsampling fMRI time series (Biswal et al., 2001), and circular bootstrap (Bellec et al., 2010, Nikolaidis et al. 2020), we expected the impact of bagging on functional connectivity estimates to exceed that of motion-ordering alone. However, performance differences between the two approaches were minimal. Since the computational demands of the motion-ordered approach are much lower than the bagging approach (motion-ordered: 6,500 correlations; bagging: 3.25 millions correlations), we recommend the motion-ordering approach as a primary data rescue approach that can enable the computation of robust and reproducible functional connectivity and brain-behavior relationships when including participants who would normally be excluded due to excess motion. Nonetheless, it is important to highlight that bagging offers advantages in other contexts. In addition to improving the reproducibility of functional brain parcellations as previously described, bagging also improves brain state classification accuracy (Richiardi et al., 2011). Participant-level resampling of training data and aggregation of model parameters and features also have been shown to improve the accuracy and generalizability of brain-behavior models for a range of phenotypic measures (e.g., fluid intelligence, sustained attention, cognitive flexibility, processing speed) using connectome-based modeling as well as other prediction frameworks (O’Connor et al., 2021; Wei et al., 2020).

### Importance of retaining high-motion participants with their usable data

While several head motion mitigation strategies have been proposed (Parkes et al., 2018; Ciric et al., 2017; Yan et al., 2013a; Yan et al., 2013b), most have focused on excluding individuals, rather than maximizing inclusion. One recent study by Smith et al. (2022) proposed a framework to enable the retention of data from first grade high-motion children (*N* = 101, 6-8 years) by employing two motion correction procedures (i.e., scrubbing and ICA-FIX) and two quality control metrics (i.e., functional connectivity and temporal signal-to-noise ratio). By varying the mean FD thresholds, the authors showed that a mean FD = 0.30 mm would be sufficient to remove most motion artifacts by retaining ∼83% of the first grade children (Smith et al., 2022). However, it is difficult to recommend an ideal threshold that would mitigate motion artifacts consistently well across different populations. The main advantage of our approaches is that they minimize the risk of discarding large amount of high-motion participants. Excluding tens, hundreds, or even thousands of these high-motion participants leads to a potentially unethical waste of participant time and effort as well as research resources, considering the time, cost, and effort invested in data collection and pre-/post-processing the imaging data, as well aso the infrastructure needed to store the raw/preprocessed data. Moreover, the lack of sample inclusivity and diversity that results from strict exclusion criteria is problematic in its own right (it has the potential to be discriminatory and compound pre-existing under-representation of younger and participants experiencing more clinically severe symptoms) and means that the estimated brain-behavior relationships may not be reproducible in other studies (Ricard et al., 2023). The motion-ordered approach may enable researchers to strike a better balance between data quality and data inclusivity.

### Limitations

Although we showed that the motion-ordered and bagging approaches can produce robust and reproducible brain-behavior associations in developing youth, the current study has several limitations. First, both approaches require the fMRI time series to be scrubbed using a head motion threshold, i.e., a proportion of volumes has to be removed from the total number of volumes acquired after being confounded by head motion. If only a small number of volumes have been acquired (e.g., 100) and/or if the head motion threshold is particularly stringent, then a large number of participants may still be excluded, due to their having an insufficient number of low-motion time points, even if the aim is to retain as few as 60 time points. Second, the motion-ordered and bagging approaches are effective mainly for functional connectivity applications that do not rely on the temporal contiguity of the fMRI time series. The reordering of motion-limited time points may undermine the assumptions of analyses that incorporate autocorrelation or other information across contiguous time points, or where the two approaches would disrupt alignment of time points across individuals, such as in intersubject correlations. Third, fingerprint ID accuracy was tested using scans collected within sessions, since these were the data available to us through the HBN dataset. Comparing the reproducibility of motion-ordered and bagged connectomes across larger spans of time would be desirable in future work. Lastly, the motion-ordered and bagging approaches were associated with a decrement in fingerprint ID accuracy relative to the standard approach, likely due to the shortened time series. There is considerable scope for future work to address these limitations and to further probe the utility, reproducibility, and validity of the motion-ordered and bagging approaches and how these should be weighed against the increased inclusivity they enable.

## 5 Conclusions

Motion-ordering can generate functional connectomes that are reproducible and which exhibit robust brain-behavior relationships in a sample of developing youth, while minimizing computational resource use. Applying motion-ordering can enable the retention of high-motion participants who would otherwise be excluded from typical brain-behavior association studies, thus maximizing participant inclusivity. Motion ordering may thereby reduce the tension between sample size requirements and data quality standards in neuroimaging, and in doing so, mitigate against resource wastage and the ethically questionable practice of discarding large numbers of participants who have given their time, effort, and trust to the research process. While the results of this study provide proof-of-concept and preliminary support for the motion-ordered approach, future work should probe its utility, reproducibility, and validity in greater depth, and across samples and phenotypes.

## Ethics

The study procedure which used the Healthy Brain Network [HBN] was approved by the School of Psychology Research Ethics Committee, Trinity College Dublin. All neuroimaging and phenotypic data were collected and openly shared in accordance with ethical approval from the Institutional Review Board, as documented here: http://fcon_1000.projects.nitrc.org/indi/cmi_healthy_brain_network/. Written consent was provided by all the participants who were at least 18 years old, while written consent was obtained from legal guardians and from the participants who were under 18 years old.

## Data and Code Availability

The analyses in this study were conducted in Python and all code is available in our GitHub repository (https://github.com/JRam02/bagging). We used publicly available packages such as pingouin (https://pingouin-stats.org/api.html), scikit-learn (https://scikit-learn.org/stable/), and SciPy (https://www.scipy.org). We also used Python libraries for data manipulation and visualization such as Numpy (https://numpy.org), Pandas (https://pandas.pydata.org), Matplotlib (https://matplotlib.org), and Seaborn (https://seaborn.pydata.org). Multiple comparison correction was performed with the False Discovery Rate (FDR) method using the statsmodels package (https://www.statsmodels.org). The areas under the curve for the standard, motion-ordered, bagging, and one-sample bootstrap approaches were computed using scikit-learn (https://scikit-learn.org/stable/modules/generated/sklearn.metrics.auc.html). The neuroimaging and phenotypic data that were used in this study are available on the HBN platform which are accessible via LORIS and COINS. We also provide the mean time series of the participants for REST1 and REST2 derived from HBN in our GitHub repository.

## Author Contribution Statement

**Jivesh Ramduny:** Conceptualization, Methodology, Formal analysis, Investigation, Writing — original draft, review & editing. **Tamara Vanderwal:** Investigation, Writing — original draft, review & editing.

**Clare Kelly:** Conceptualization, Methodology, Formal analysis, Investigation, Writing — original draft, review & editing, Funding, Supervision, Project administration.

## Acknowledgment

We thank the Healthy Brain Network for allowing the neuroimaging and phenotypic data to be openly accessible via LORIS and COINS.

