## Supplementary Materials for "Data rescue in high-motion youth cohorts for robust and reproducible brain-behavior relationships"

### Supplementary Material

#### Influence of minTP and TP on bagged brain-behavior relationships

Supplementary Figure 1 illustrates the influence of minTP (minTP ϵ {60, 100, 200}) and (minTP ϵ {20, 60, 100}). Varying the minTP threshold revealed a non-linear relationship between the widths of the 95% CIs and sample size. Across the three minTP thresholds, the widths of the 95% CIs were considerably large ($width_{minTP60}$ = 0.78; $width_{minTP100}$ = 0.82; $width_{minTP200}$ = 0.77) at typical sample sizes (*N* = 20) but narrowed non-linearly (width = 0.01 for each minTP) with increasing number of participants (*N* = 255). When we varied the TP thresholds, we detected a similar non-linear relationship between the widths of the 95% CIs and sample size. Across the three TP thresholds, the widths of the 95% CIs were large ($width_{TP20}$ = 0.78; $width_{TP60}$ = 0.84; $width_{TP100}$ = 0.83) at small sample sizes (*N* = 20) but non-linearly tightened (width = 0.01 for each TP) as the sample size became large (*N* = 255).

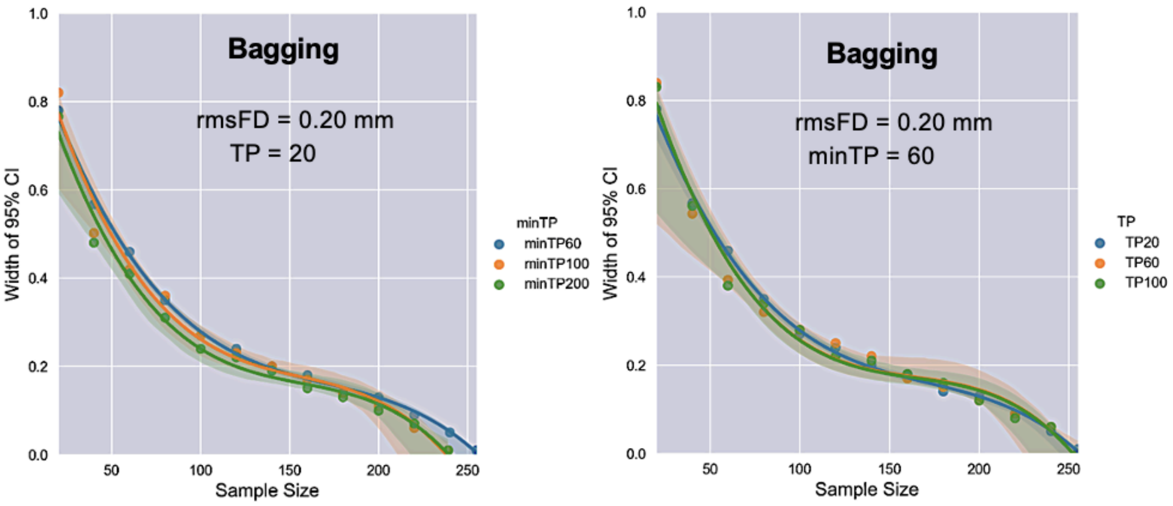


**Supplementary Figure 1.** (A) Influence of minTP using bagging on the width of the 95% CIs by keeping TP constant at 20 such that minTP ϵ {60, 100, 200}. (B) Influence of TP using bagging on the width of the 95% CIs by keeping minTP constant at 60 such that TP ϵ {20, 60, 100}.

**Characteristics of low-motion (*N* = 255) and "rescued" high-motion (*N* = 124) participants**

Relative to the low-motion participants, high-motion participants were twice as likely to be male (two-way Chi-Square test, $\mathcal{X}^2 = 6.93$, $P = 0.008$; odds ratio of male to female participants is 1.9), exhibited greater average motion [rmsFD] for REST1 (Mann-Whitney U test, U = 1,870.5, $P < 0.001$) and REST2 (Mann-Whitney U test, U = 450.5, $P < 0.001$). There were also significant differences in the distributions for REST1 rmsFD (two-sample Kolmogorov Smirnov (KS) test, D = 0.75, $P < 0.001$), REST2 rmsFD (two-sample KS test, D = 0.88, $P < 0.001$), and CBCL-internalizing (two-sample KS test, D = 0.16, $P < 0.023$) when comparing the two groups (**Supplementary Figures 2-3**).

On average, the "rescued" high-motion participants were younger (mean age = 9.8 years) relative to the low-motion participants (mean age = 11.8 years; Mann-Whitney U test, U = 21,878.5, $P < 0.001$). The age distributions for the two groups were also statistically different (two-sample Kolmogorov Smirnov test, D = 0.40, $P < 0.001$).

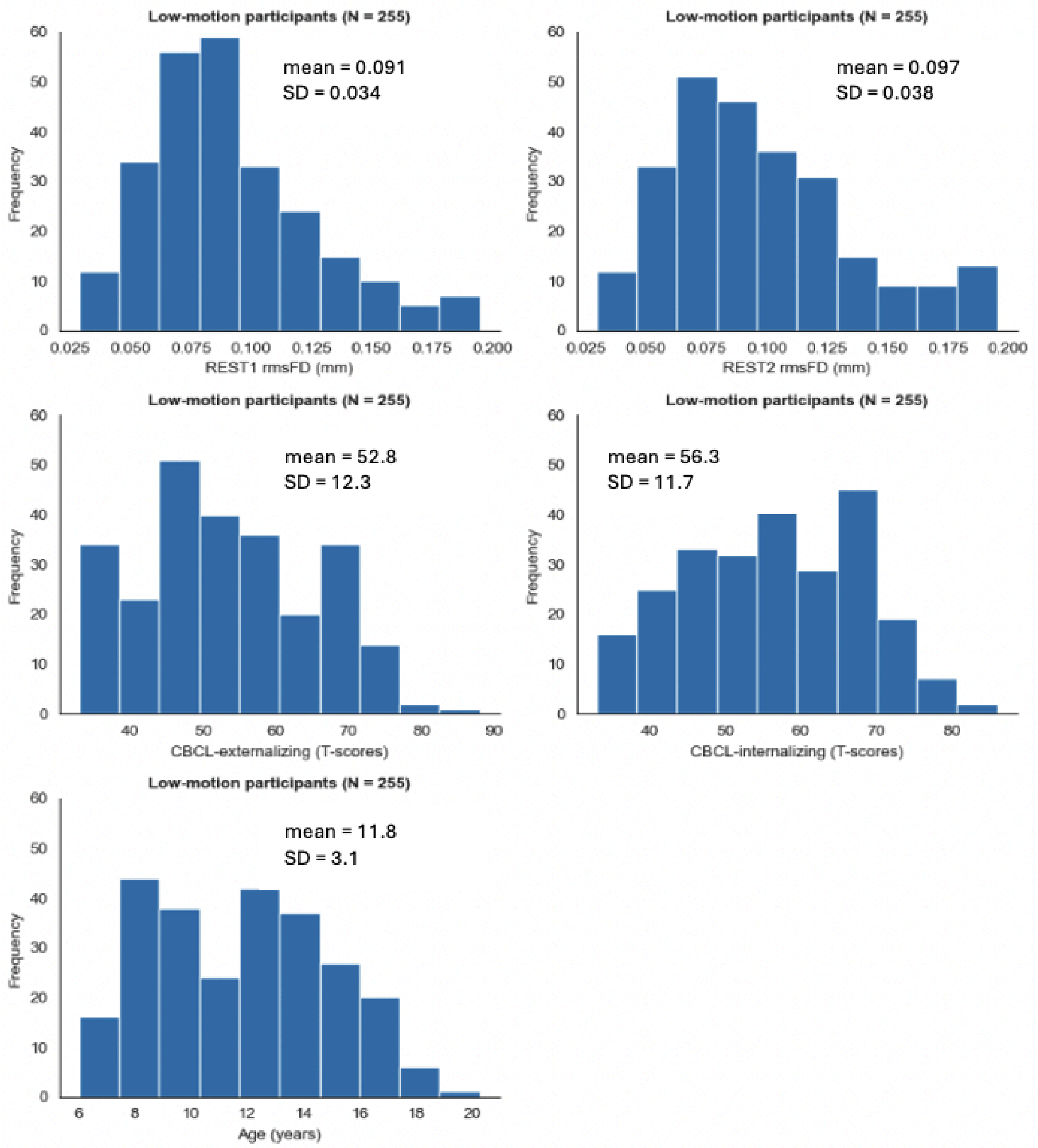


**Supplementary Figure 2.** Demographic, behavioral, and imaging characteristics of the low-motion participants (*N* = 255). The externalizing and internalizing behaviors of the participants are measured by their respective subscales from the Child Behavior Checklist [CBCL]. The CBCL scores are T-standardized. The mean and standard deviation for each characteristic is also displayed.

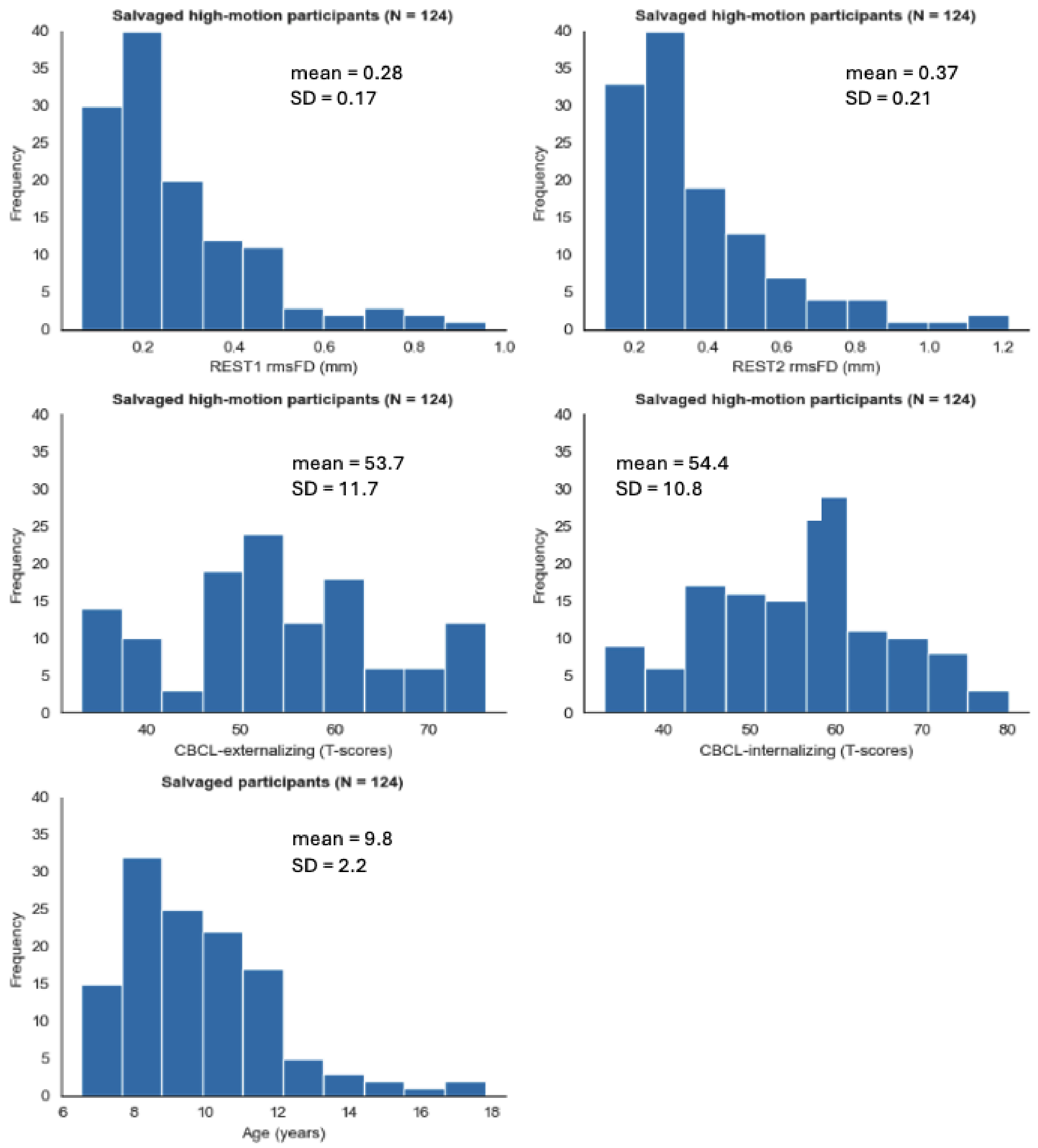


**Supplementary Figure 3.** Demographic, behavioral, and imaging characteristics of the “rescued” high-motion participants (*N* = 124). The externalizing and internalizing behaviors of the participants are measured by their respective subscales from the Child Behavior Checklist [CBCL]. The CBCL scores are T-standardized. The mean and standard deviation for each characteristic is also displayed.
